# Unifying ecosystem resistance, resilience, and recovery from extreme stress into a single statistical framework

**DOI:** 10.1101/743708

**Authors:** Nathan P. Lemoine

**Affiliations:** Department of Biological Sciences, Marquette University, Milwaukee, WI 53233; Department of Zoology, Milwaukee Public Museum, Milwaukee, WI 53233

## Abstract

Natural communities and ecosystems are currently experiencing unprecedented rates of environmental and biotic change. While gradual shifts in average conditions, such as rising mean air temperatures, can significantly alter ecosystem function, ecologists recently acknowledged that the most damaging consequences of global change will probably emanate from both a higher prevalence and increased intensity of extreme climatic stress events. Given the potential ecological and societal ramifications of more frequent disturbances, it is imperative that we identify which ecosystems are most vulnerable to global change by accurately quantifying ecosystem responses to extreme stress. Unfortunately, the lack of a standardized method for estimating ecosystem sensitivity to drought makes drawing general conclusions difficult. There is a need for estimates of resistance/resilience/legacy effects that are free of observation error, not biased by stochasticity in production or rainfall, and standardizes stress magnitude among many disparate ecosystems relative to normal interannual variability. Here, I propose a statistical framework that estimates all three components of ecosystem response to stress using standardized language (resistance, resilience, recovery, and legacy effects) while resolving all of the issues described above. Coupling autoregressive time series with exogenous predictors (ARX) models with impulse response functions (IRFs) allows researchers to statistically subject all ecosystems to similar levels of stress, estimate legacy effects, and obtain a standardized estimate of ecosystem resistance and resilience to drought free from observation error and stochastic processes inherent in raw data. This method will enable researchers to rigorously compare resistance and resilience among locations using long-term time series, thereby improving our knowledge of ecosystem responses to extreme stress.

## Introduction

Natural communities and ecosystems are currently experiencing unprecedented rates of environmental and biotic change. While gradual shifts in average conditions, such as rising mean air temperatures, can significantly alter ecosystem function, ecologists recently acknowledged that the most damaging consequences of global change will probably emanate from both a higher prevalence and increased intensity of extreme climatic stress events, defined as climate events that occur outside the statistical bounds of historical conditions (Smith 2011). For example, more frequent and severe droughts in North America and Europe have already caused shifts in plant community composition, widespread tree mortality, and catastrophic declines in primary production (Ciais et al. 2005, Anderegg et al. 2013, Hoover et al. 2014, Knapp et al. 2015a). Concurrent with drought, the frequency and duration of heat waves have also increased over the past century (Perkins et al. 2012). In terrestrial systems, heat waves exacerbate drought water stress by increasing evapotranspiration, while marine heat waves cause extensive mortality of foundation species and habitat loss (Ciais et al. 2005, Le Nohaïc et al. 2017, Smale et al. 2019). Given the potential ecological and societal ramifications of more frequent disturbances, it is imperative that we identify which ecosystems are most vulnerable to global change by accurately quantifying ecosystem responses to extreme stress.

Ecosystem stress response consists of two components: the decline in ecosystem function during or immediately following the stress event, and the degree of improvement in ecosystem function after alleviation of stress (Lloret et al. 2011, Smith 2011). The magnitude of decline in ecosystem function during stress, often called ‘resistance’ or ‘sensitivity’ to stress, has been studied extensively and varies both among and within ecosystems (Huxman et al. 2004, Knapp et al. 2015, Sully et al. 2019). Grasslands, for instance, are typically more drought-sensitive than forests because grasslands occupy drier climate conditions (Stuart-Haëntjens et al. 2018). Indeed, arid and semi-arid grasslands are among the most drought-sensitive ecosystems on the planet, losing up to twice as much primary production, proportionally, than mesic grasslands during drought (Knapp et al. 2015a). In marine ecosystems, coral reef resistance to bleaching also depends on local climate conditions. Reefs in thermally variable water bodies bleach less extensively than reefs in thermally constant environments (Sully et al. 2019). Thus, a relatively large body of literature exists to describe how ecosystem sensitivity to extreme climatic stress varies with either abiotic or biotic variables. In comparison, the abiotic and biotic processes that accelerate or inhibit restoration of ecosystem function after climate stress remain poorly understood. The consequences of extreme stress can persist for some time following perturbation (*i.e.* legacy effects; Smith 2011, Sala et al. 2012, Anderegg et al. 2015), but few multi-site studies have assessed the climatic and biological determinants of ecosystem recovery. Furthermore, it is difficult to infer general patterns and mechanisms regarding differential ecosystem sensitivity to and recovery from climate stress because there is no standardized vocabulary describing ecosystem responses to stress, and current quantitative method for estimating ecosystem trajectories during and after stress suffer from significant statistical problems.

Consider the issue of estimating resistance/sensitivity of terrestrial primary production to drought. The first problem is the lack of a consistent method for describing the decline in primary production; resistance and sensitivity are antonyms for the same phenomenon with at least five common mathematical formulations (Table 1). The second problem is that comparing sensitivity/resistance estimates among sites or years is difficult even when using a single metric. For example, estimating sensitivity/resistance as the ratio of production during a drought year to production during the previous year (Table 1) biases estimates of sensitivity/resistance if the year prior to drought was below or above average rainfall. That is, the context of the drought varies from site to site and year to year, impairing inter-site or even intra-site comparisons of drought sensitivity. This ratio also includes observation error within its estimate; low or high estimates of primary production during either the previous year or the drought year partly arise from imperfect sampling methods, such that we cannot know the extent to which the ratio represents the true process of resistance or incorporates sampling artifacts. Finally, it is difficult to place estimates of sensitivity into a climate context because the degree of stress induced by a given rainfall reduction varies among grasslands. That is, a 200mm reduction in annual rainfall imposes a much stronger meteorological drought in the arid shortgrass steppe than it does in mesic tallgrass prairies (Knapp et al. 2015b). Thus, there is a need for an estimate of sensitivity/resistance that is free of observation error, not biased by stochasticity in production or rainfall, and standardizes stress magnitude among many disparate ecosystems relative to normal interannual variability.

**Table 1.** Definitions and mathematical equations used to calculate ecosystem resistance, resilience, recovery, and legacy effects following an extreme stress event.

| Method | Name | Equation | Units | Citation |
| --- | --- | --- | --- | --- |
| **Reduction During Stress** |  |  |  |  |
| 1 | Sensitivity | $\frac{x_t - x_{t-1}}{ppt_t - ppt_{t-1}}$ | Change in primary production per mm chain in rainfall | (Wilcox et al. 2017) |
| 2 | Sensitivity | $\frac{\Delta x}{\Delta ppt}$ | Slope of the primary production – precipitation relationship | (Huxman et al. 2004, Knapp et al. 2015a) |
| 3 | Sensitivity | $100 \times \left( \frac{x_t - \bar{x}}{\bar{x}} \right)$ | Percent decline from long-term mean | (Griffin-Nolan et al. 2018) |
| 4 | Resistance | $\frac{x_t}{x_{t-1}}$ | Proportion decline from pre-drought year | (Lloret et al. 2011, Gazol et al. 2017, 2018) |
| 5 | Resistance | $\ln \left( \frac{x_t}{x_{t-1}} \right)$ | Log proportion decline from pre-drought year | (Stuart-Haëntjens et al. 2018) |
| **Return Following Stress** |  |  |  |  |
| 6 | Recovery | $\frac{x_{t+1}}{x_t}$ | Proportion increase in post-drought year | (Lloret et al. 2011, Gazol et al. 2017, 2018) |
| 7 | Resilience | $\frac{x_{t+1}}{x_{t-1}}$ | Proportion decrease in post-drought year from pre-drought | (Lloret et al. 2011) |
| 8 | Resilience | $\ln \left( \frac{x_{t+1}}{x_{t-1}} \right)$ | Log proportion decrease in post-drought year from pre-drought | (Stuart-Haëntjens et al. 2018) |
| 9 | Legacy effects | $100 \times \left( \frac{x_{t_1} - \bar{x}}{\bar{x}} \right)$ | Percent decrease in post-drought year from long-term mean | (Griffin-Nolan et al. 2018) |
| 10 | Legacy Effects | $x_{t+1} - \hat{x}_{t+1}$ | Observed – predicted for post-drought year | (Sala et al. 2012, Anderegg et al. 2015) |

Quantifying ecosystem resilience/recovery/legacy effects to climate stress has proven no less idiosyncratic (Table 1), and many of the same statistical artifacts described above afflict widely used estimates of resilience/recovery. As before, estimating ecosystem resilience using the proportion reduction in primary production after drought compared to primary production before drought incorporates both temporal stochasticity and observation error. Resilience to drought might, for example, be overestimated if the post-drought year is abnormally wet or the pre-drought year was abnormally dry, or if observation error resulted in higher measurements of primary production post-drought simply due to sampling protocols. Perhaps the most popular method for quantifying resilience/legacy effects of primary production is to calculate the predicted primary production for every year using a temporal primary production-precipitation regression based on inter-annual time series data (Sala et al. 2012, Anderegg et al. 2015). The error of the post-drought year, observed – predicted, constitutes the legacy effect. To demonstrate, the shortgrass steppe of Colorado experienced an extreme drought in 2012 (Fig. 1A – red dot). Based on the primary production-precipitation relationship, we can estimate the predicted value of primary production in 2013 (Fig. 1B). The legacy effect of drought is then the observed primary production (Figs. 1A,B – green dot) minus the predicted primary production in 2013 (Fig. 1B).

**Figure 1.**
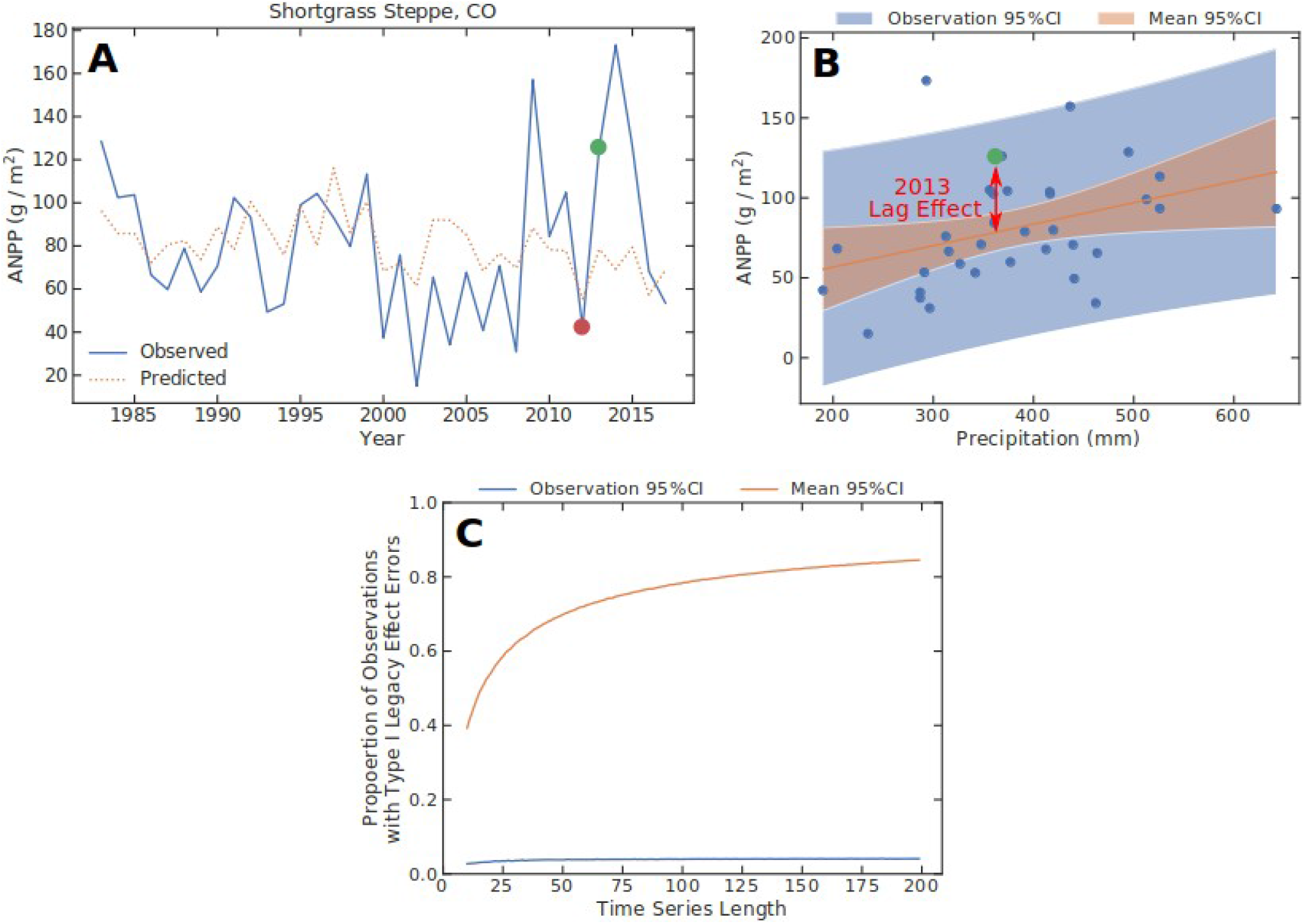
**A)** Time series of aboveground net primary production at the shortgrass steppe near Fort Collins, Colorado. The red point denotes the drought of 2012, the green point denotes primary production in the year following drought. The orange dashed line shows predicted primary production based on the primary production – precipitation relationship. **B)** The primary production – precipitation relationship for the shortgrass steppe. The orange shaded area is the 95% CI of the mean, while the blue shaded area is the 95% CI of individual observations. Green point shows the recovery year of 2013 and how “legacy effects” are sometimes calculated (Table 1). **C)** Using the mean 95% CI (orange line) to statistically test for legacy effects results in high false positive rates as sample size increases and uncertainty about the mean decreases, while using the observation 95% CI (blue line) avoids this complication. Lines were generated by simulating 10,000 precipitation time series, then using a simulating primary production – precipitation relationship to estimate primary production in the absence of legacy effects. Type I error rates are the proportion of observations in a simulated time series that would be considered to possess significant legacy effects, despite being simulated without legacy effects.

However, this estimation method suffers from several logical and statistical issues. First, observations will almost never fall exactly along the regression line; by definition half of the points will be above and half will be below the line. Second, the distance from the regression line that we consider to be a significant ‘legacy’ effect depends generally on sample size and less on any biological property. Third, this method assumes that scatter surrounding the primary production-precipitation relationship is entirely caused by legacy effects. Is this true for all points, or only for the one point in which the ecologist is interested? If legacy effects are the only cause of scatter, then incorporating legacy effects into regression models should perfectly model data. Though autoregressive parameters sometimes do improve fit, they do not model data perfectly nor do they substantially improve prediction accuracy for a single observation (Oesterheld et al. 2001). If legacy effects apply only to the year following an extreme event, do sources of variability present in other years not occur in post-stress years, such that legacy effects explain the entirety of deviation from the mean in the post-stress year? In reality, many factors likely contribute to an imperfect primary production-precipitation relationship in all years, including the within-year distribution of rainfall event size and timing (Heisler-White et al. 2008), observation error, stochasticity in community composition, and potential legacy effects. Sites with a weaker primary production-precipitation relationship (*i.e.* lower *R*^2^) will have more scatter about the line and therefore possess stronger “legacy effects”. There is currently no way to parse out whether legacy effects calculated in this manner arise from true legacy effects, observation error, or how the legacy effect depends on the actual weather patterns during the recovery year.

To impose statistical rigor in testing for legacy effects, some studies compared the observed point to the 95% CI of the predicted value and report only legacy effects that are significantly different from the mean (Griffin-Nolan et al. 2018) (Table 1; Fig. 1B - orange shaded area). However, this too has statistical issues. First, as above, the presence of a point inside or outside the mean CI might be observation error. Second, the significance of a legacy effect depends on the width of the mean 95% CI, which is at least partially determined by sample size. For long time series, over 80% of observations would be considered significant legacy effects when compared to the mean CI, even when data were generated without legacies (Fig. 1C). In other words, sites with longer time series (*i.e.* more data) are more likely to show significant “legacy effects” even when no legacy effects are present because the width of the 95% CI shrinks proportionally to the inverse square-root of sample size. Comparing the presence of legacy effects among sites might simply be comparing differences in time series length. In the shortgrass steppe, the point for 2013 would be a ‘significant’ legacy effect even though it falls well within the 95% envelope for individual data points (Fig. 1B). To test whether legacy effects fall outside the normal range of variability, it is more appropriate to compare the observed ‘legacy effect’ point to the observation/prediction interval. Using the observation confidence interval, instead of the mean confidence interval, alleviates this particular issue by minimizing false legacy effects because the width of the observation interval depends only on residual error, not sample size (*i.e.* Type I error for simulations, Fig. 1C). This issue raises the possibility that many reported legacy effects might be noise, and ecosystem ecologists currently have no statistical method that can reliably separate legacy effects from observation error.

Here, I propose a statistical framework that estimates all three components of ecosystem response to stress using standardized language (resistance, resilience, recovery, and legacy effects) while resolving all of the issues described above. Coupling autoregressive time series with exogenous predictors (ARX) models with impulse response functions (IRFs) allows researchers to statistically subject all ecosystems to similar levels of stress, estimate legacy effects, and obtain a standardized estimate of ecosystem resistance and resilience to drought free from observation error and stochastic processes inherent in raw data. This method will enable researchers to rigorously compare resistance and resilience among locations using long-term time series, thereby improving our knowledge of ecosystem responses to extreme stress.

### Impulse Response Functions

Impulse response functions derive from time series analyses (*e.g.* autoregressive and/or moving average models) and describe the trajectory of dynamic systems following stress. They are particularly useful in systems that are logistically difficult, costly, or impossible to manipulate experimentally, such as financial markets. Indeed, econometricians have widely implemented IRFs to understand the resistance, resilience, and recovery of financial markets to instantaneous “shocks” (Creal and Wu 2017, Gambetti and Musso 2017). For example, Senbet (2016) used IRFs to visualize the consequences of higher federal interest rates on unemployment, consumption, and other indicators of economic health. In medical studies, IRFs describe how the human body responds to pulsed stress events, such as elevated or depressed hormone activity (Schultz et al. 2015, Chang et al. 2017). Earth system modelers use IRFs to understand how global temperature or CO_2_ concentrations respond to various shocks, such as changes in oceanographic processes or vehicular emissions (Thompson and Randerson 1999, Joos et al. 2013, Millar et al. 2017, Zeng et al. 2017). However, no ecological study has yet used IRFs to quantify ecosystem resistance to and recovery from extreme stress events. Fortunately, calculating IRFs is as simple as fitting autoregressive models using standard time series methods readily available in many programming languages (*e.g.* the arimax function in the TSA library of R), comparing models to determine the appropriate autoregressive order, and then calculating the components of ecosystem stress response using simple combinations of parameters from the best model. The ease of IRF calculation could facilitate widespread adoption in assessing ecosystem responses to extreme stress across a variety of study systems.

Constructing IRFs first requires fitting various ARX(*p*) models to long-term time series data to identify whether legacy effects are present. ARX(*p*) models modify autoregressive models of order *p* (*i.e.* AR(*p*) models) by including one or more exogenous variables:

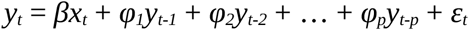

This model states that ecosystem state at time *t* (*y*_*t*_) depends on current exogenous values (*x*_*t*_, *e.g.* annual precipitation, sea-surface temperature anomaly, etc.), previous ecosystem states up to *p* time steps in the past (*y*_*t−p*_, *i.e.* legacy effects), and error from both unmeasured processes and sampling issues (*ε*_*t*_). The appropriate order *p* can be chosen via information theoretic methods (*e.g.* AIC, BIC) or via chi-square likelihood ratio tests comparing successively lower orders (*e.g.* AR(2) vs. AR(1), AR(1) vs. AR(0), etc). The lowest order model, ARX(0), is simply a linear regression of ecosystem state against the exogenous variable with no intercept if the response data have been standardized prior to regression (the intercept is the mean, and standardization of the response makes the mean equal to 0). Both *y* and *x* should be standardized to a mean of 0 and standard deviation of 1, especially if the objective is to compare stress resistance and resilience among different study sites or ecosystems.

Once the appropriate ARX(*p*) model has been identified, the next step is to derive the IRF. IRFs use the fitted ARX(*p*) model to predict ecosystem state under new values of *x* through time. The special case of imposing an initial stress to the exogenous variable (*e.g.* drought or heat wave) then allowing the exogenous variable to return to nominal levels for recovery is represented by:

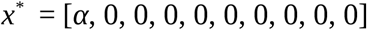

where *α* denotes the level of stress at the initial time point. It is critically important that *x* be standardized prior to model fitting, thus a value of *x*^*^_*t*_ = 0 represents average exogenous conditions (*i.e.* average precipitation) and *α* is stress in units of standard deviations for the exogenous predictor. In this way, ecologists can statistically subject disparate ecosystems to the same level of relative stress (e.g. a 2*σ* decline in precipitation) and allow the system to return to nominal values when estimating ecosystem resistance and resilience to stress, thereby eliminating stochasticity and observation error present in current methods.

After determining the appropriate value for *α*, the IRF for ecosystem state (*y*^*^) can be calculated either recursively or, in the case of the simple *x*^*^ used here, analytically. Ecologists can then use the IRF to quantify various components of ecosystem response to stress. In the context of IRFs, stress responses can be defined as:

1. **Resistance –** The standardized decline in ecosystem state when subjected to an initial stress of *α* in the exogenous variable at *t* = 0. More negative values imply lower resistance. For example, stronger drought-induced declines in primary production, or bleaching-induced reductions in coral cover, yield more negative resistance values. Units are in standard deviations of the ecosystem state, provided that *y* has been appropriately standardized beforehand.
2. **Resilience**– The standardized decline in ecosystem state in the time step immediately following an initial stress of *α* in the exogenous variable. More negative values imply lower resilience. Positive values indicate positive legacy effects. Such situations occur when, for example, drought causes a buildup of soil nitrogen that can stimulate plant growth when rainfall returns to normal levels the following year (Hofer et al. 2017). Units are in standard deviations of the ecosystem state, provided that *y* was appropriately standardized beforehand.
3. **Recovery –** The amount of time required for the ecosystem state to recover to half of its resistance value. In other words, how much time is required for the ecosystem state to regain a given percentage of the decline experienced during stress. Larger values imply slower recovery. Units are in time steps *t* (*i.e.* years for annual primary production). Note that I in this paper, I chose 50% recovery arbitrarily and ecologists could modify this value depending on the question or system under study.

For the simplified *x*^*^ listed above, each quantity has analytical solutions for ARX(*p*) models of different orders (Table 2).

**Table 2.**
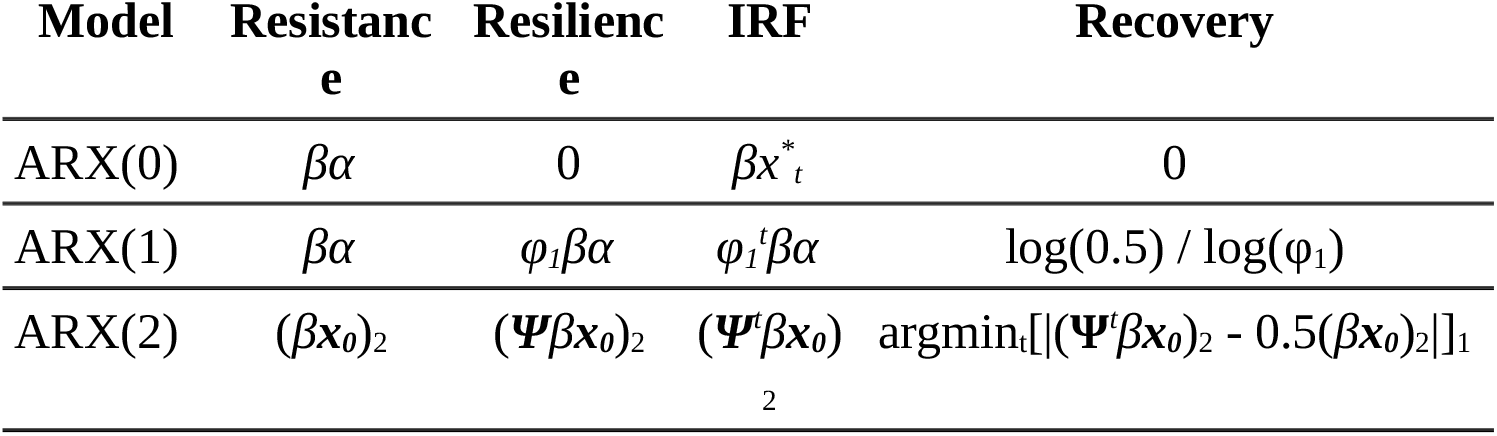
Analytical equations for calculating resistance, resilience, IRF, and recovery for ARX models of different orders. Note that these equations rely on the special form of *x*^*^ described in text. More complex solutions exist for general forms of *x*^*^. The subscript _2_ denotes the second element of a column vector (see text).

#### ARX(0) Models

The ARX(0) model is a simply linear regression of standardized ecosystem state against a standardized exogenous variable. The IRF is given by:

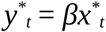

Resistance to the initial stress at *t*^*^ = 0 is then:

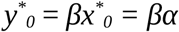

In this model, there are no ecosystem resilience or recovery values because the ecosystem recovers immediately and perfectly in the absence of legacy effects. That is, the ecosystem state returns to average conditions following the return of the exogenous value to average conditions:

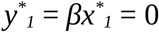

#### ARX(1) Models

ARX models with one lag effect are given by the equation:

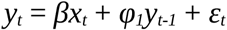

In this case, the IRF can be calculated via recursive substitution:

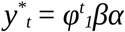

Assuming that the ecosystem state was at average conditions prior to stress (*y*^*^_*−1*_ = 0) this gives ecosystem resistance as:

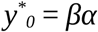

Resilience is ecosystem state at the next time step:

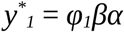

Recovery is the time required to achieve a 50% return to nominal levels:

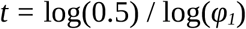

#### ARX(2) Models

ARX(2) models include two lagged time steps:

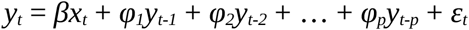

General solutions require linear algebra. The equation above can be placed into matrix algebra form as:

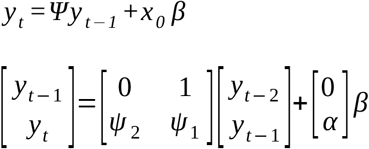

which, when expanded, yields the two equations:

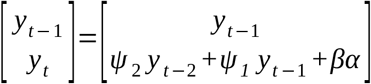

Assuming that both the ecosystem state and exogenous values were at average conditions prior to the stress (*i.e. y*^*^_*−2*_ = *y*^*^_*−1*_ = 0), then resistance is given by:

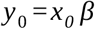

However, because this is a vector of length 2, resistance at *t* = 0 is actually the second element of the vector:

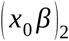

Recursive substitution yields the value for resilience at *t* = 1:

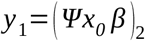

where again the subscript denotes the second element of the vector. More generally, the IRF has the form:

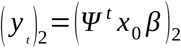

Importantly, ***Ψ***^*t*^ is the matrix power of *t*^*1*^. Recovery cannot be calculated analytically because ARX(2) models exhibit oscillations, and there may be more than one time point where *y*^*^_*t*_ = 0.5 *y*^*^_*0*_. As a result, recovery is best quantified as the first time point where *y*^*^_*t*_ is closest to 0.5:

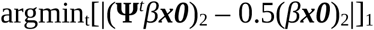

where the subscript 1 denotes the first element of the set in the event that multiple points satisfy the condition within the brackets. The interior of the brackets denotes the absolute value of the difference between the IRF and half of resistance.

### Resistance and resilience of a cool-season grassland to drought

The simplicity of IRFs can be demonstrated with an example application. I estimated the resistance, resilience, and recovery of aboveground net primary production (ANPP) in a northern, cool-grass prairie to extreme drought using primary production and precipitation data from Manyberries, Alberta (Smoliak 1986) (Figure 2). Prior to analyses, both ANPP and precipitation were examined for gap years (*n* = 4 non-sequential gap years). Because ARX models cannot work with missing years, I first interpolated the four missing production and precipitation values using a radial basis function. After standardizing ANPP and precipitation as described above, I detrended both time series by calculating the residuals of each time series regressed against time to ensure weak stationarity required by autoregressive models. I then calculated resistance and resilience following the methods in Table 1. Next, I used IRFs to estimate resistance, resilience, and recovery. To do so, I fit ARX(0), ARX(1), and ARX(2) models to the dataset using dedicated time series methods (the ARIMAX function in the *statsmodels.tsa* Python module). After model fitting, I identified the best-fitting model using Bayesian Information Criteria (BIC). I chose BIC because BIC penalizes additional terms more heavily than AIC and is therefore more conservative.

**Figure 2.**
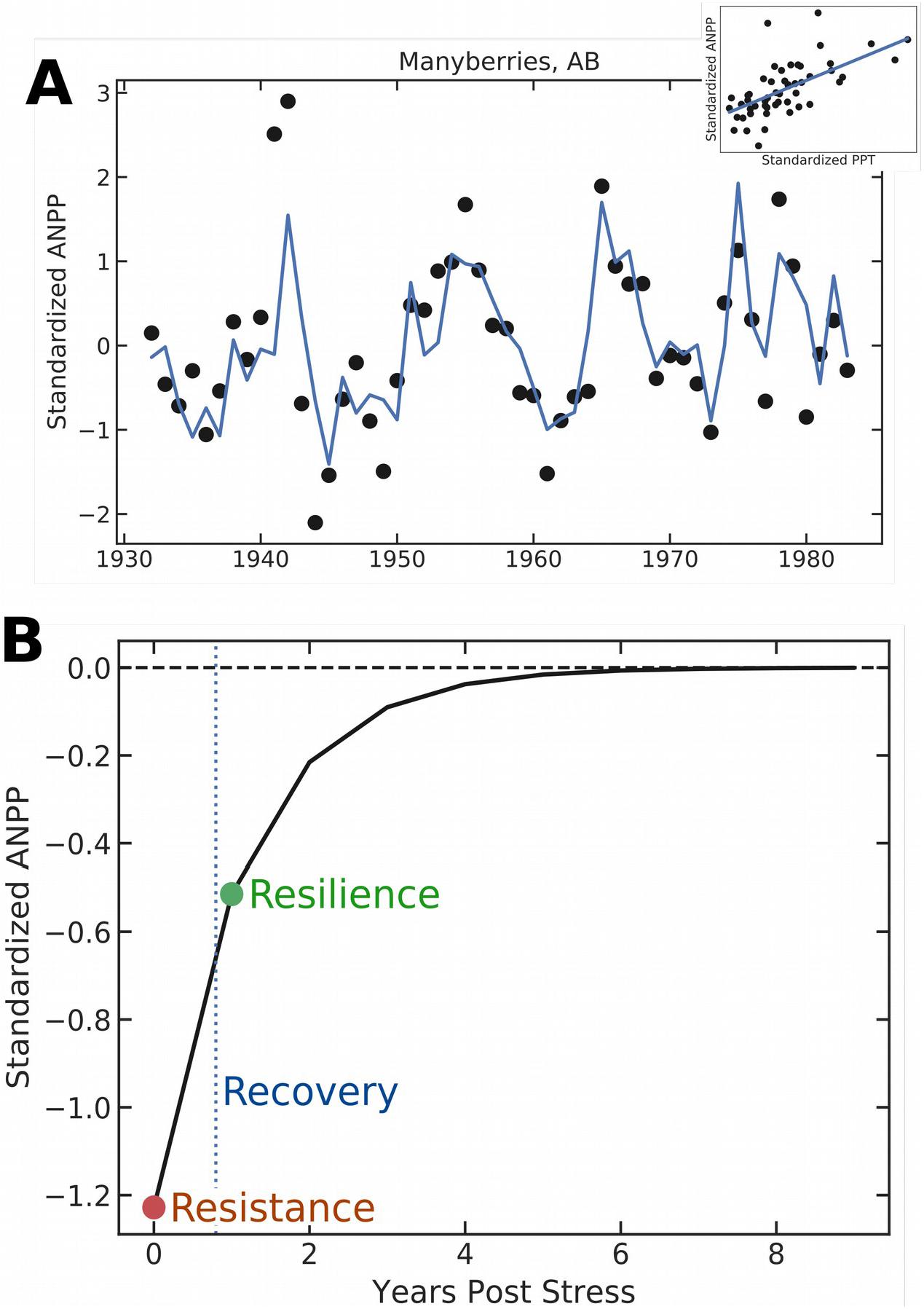
**A)** Time series of aboveground net primary production (ANPP) at Manyberries, AB. Blue line shows the fitted ARX(1) model. Inset shows the ANPP – precipitation relationship. **B)** IRF for the ARX(1) model fitted to Manyberries, AB. Values for resistance, resilience, and recovery to a 2*σ* drought were calculated following Table 2.

Primary production at Manyberries was highly contingent on annual precipitation (Figure 2A). Using Method 1 (see Table 1), the slope of the primary production – precipitation relationship provides an estimate of 0.14 ± 0.02 g ANPP per mm ppt. However, calculating sensitivity following Method 2 provides an estimate of 0.10 g ANPP per mm ppt, significantly lower than the estimate of sensitivity from Method 1 (*p* = 0.046). I then estimated drought resistance using Methods 3 and 4 for every year in which precipitation fell beneath the 5% quantile (standardized precipitation < −1.64). In Manyberries, only the year 1973 met this criteria. According to Method 3, during 1973 Manyberries experienced a 51% decline in ANPP relative to the long-term mean. However, using percent change from the previous year provided only a 36% decline in ANPP, despite the previous year possessing near average precipitation (standardized precipitation for 1972 = −0.26). This highlights the influence of potential observation error; the preceding year with average precipitation had below average ANPP due to either process or observation errors, but it is impossible to know how severely those errors biased sensitivity estimates. Thus, each method outlined in Table 1 provides significantly different, sometimes vastly so, estimates of drought sensitivity at Manyberries AB, and often on different, non-comparable scales. Furthermore, these methods cannot disentangle observation error from process error, nor incorporate potential legacy effects that might affect the prior year’s ANPP.

IRFs can rectify these issues by using ARX models. For example, BIC model selection identified clear legacy effects (Table 3). Because both one and two-year lags were equally plausible (Table 3), I chose the ARX(1) model as the most parsimonious fit to the data. Positive autocorrelation (*φ*_*1*_ = 0.42 ± 0.09) suggests that extremely low ANPP in a given year would be followed by lower than expected ANPP the next, such that drought might negatively impact ANPP for several years after the initial event. To visually and empirically quantify ecosystem responses to drought, I calculated the IRF, resistance, resilience, and recovery time of ANPP after a 2*σ* decline in rainfall (Fig. 2B). A 2*σ* drought resulted in a 1.2*σ* decline in ANPP, and strong legacy effects inhibited complete recovery for approximately 7 years (Fig. 2B). However, Manyberries production was quite resilient, having recovered 50% of its function in 0.80 years, such that the year following drought saw only a 0.51*σ* decline in ANPP (Figure 2B).

**Table 3.** BIC model selection criteria for ARX(0), ARX(1), and ARX(2) models fit to primary production/precipitation data from Manyberries, AB.

| Model | BIC | ΔBIC | Parameters<br>(mean ± 1 SE) |
| --- | --- | --- | --- |
| ARX(0) | 123.4 | 14.8 | $\beta$: 0.65 ± 0.11 |
| **ARX(1)** | **108.6** | **0.0** | $\beta$: 0.61 ± 0.09<br>$\varphi_1$: 0.42 ± 0.09 |
| ARX(2) | 109.1 | 0.5 | $\beta$: 0.58 ± 0.09<br>$\varphi_1$: 0.51 ± 0.10<br>$\varphi_2$: -0.19 ± 0.10 |

### Climate constraints on grassland resistance and resilience to drought

By standardizing both the severity of stress and ecosystem response, IRFs can be used to compare ecosystem resistance and resilience across sites of differing biotic and abiotic contexts. For example, ecologists commonly use long term time series data to assess how grasslands vary in drought resistance and resilience across global or continental climate gradients (Sala et al. 2012, Knapp et al. 2015a). However, previous efforts have been, quite understandably, hindered by the difficulties in standardizing drought effects across sites and accurately quantifying drought resistance free from temporal stochasticity. Here, I used IRFs to calculate ecosystem resistance, resilience, and recovery of 14 globally distributed herbaceous sites previously identified to possess significant legacy effects.

Using (Sala et al. 2012), I identified 14 datasets composed of both annual precipitation and aboveground primary production in herbaceous communities. Gap years in either primary production or precipitation were filled using a radial basis function. Ideally, time series would contain at least thirty consecutive years of data. Unfortunately, very few ecological datasets span that duration. I therefore kept datasets with ten or more years. Prior to analyses, I gap-filled, standardized, and detrended each dataset as was done for Manyberries, AB (see above). To illustrate the disparity among existing methods for estimating ecosystem stress resistance, I calculated resistance for each dataset using two ‘slope-based’ methods (Methods 1 and 2, Table 1) and two ‘percent-based’ methods (Methods 3 and 4, Table 1). I then used BIC to distinguish between ARX(0), ARX(1), and ARX(2) models. After identifying the appropriate autoregressive order, I calculated IRFs, resistance, resilience, and recovery following a 2*σ* decline in precipitation. In this way, ecosystem resistance, resilience, and recovery were all derived for the same magnitude of rainfall reduction relative to ambient conditions at each site. To assess climate constraints on ecosystem resistance, resilience, and recovery from drought, I regressed each metric against mean annual precipitation derived from WorldClim.

The four resistance methods gave quantitatively different results. The two ‘slope-based’ methods were poorly correlated (*r* = 0.24), largely due to the presence of one outlier from Method 2 (Fig. 3A). The outlier exemplifies the danger of Method 2, representing a site with a large change in ANPP but little change in precipitation between pre-drought to drought years. The degree to which this point is influenced simply by observation error between years is unknown. The remaining points were scattered around the 1:1 line but, importantly, the order of sites differed between Methods 1 and 2 (Fig. 3A). The least sensitive site identified by Method 1 was only the fourth least resistant site in Method 2, while the most sensitive site according to Method 1 was only the fifth most sensitive site according to Method 2 (Fig. 3A). Re-ordering issues were less severe between the two ‘percent-based’ methods, largely due to the higher degree of correlation between Methods 3 and 4 (*r* = 0.83, Fig. 3B). However, comparing across ‘slope-based’ and ‘percent-based’ methods yielded much weaker correlation among sensitivity estimates (Methods 1 vs. 3: *r* = 0.47, Fig. 3C; Methods 1 vs. 4: *r* = −0.42, Fig. 3D) and, as a result, strong reordering of sites. For example, the most sensitive site identified by Method 1 was the fourth-most sensitive site in Method 3, while the most sensitive site in Method 3 was ninth-most sensitive site in Method 1. Thus, there is a strong need for a method of estimating sensitivity that is applied consistently across studies to facilitate comparisons across different time series.

**Figure 3.**
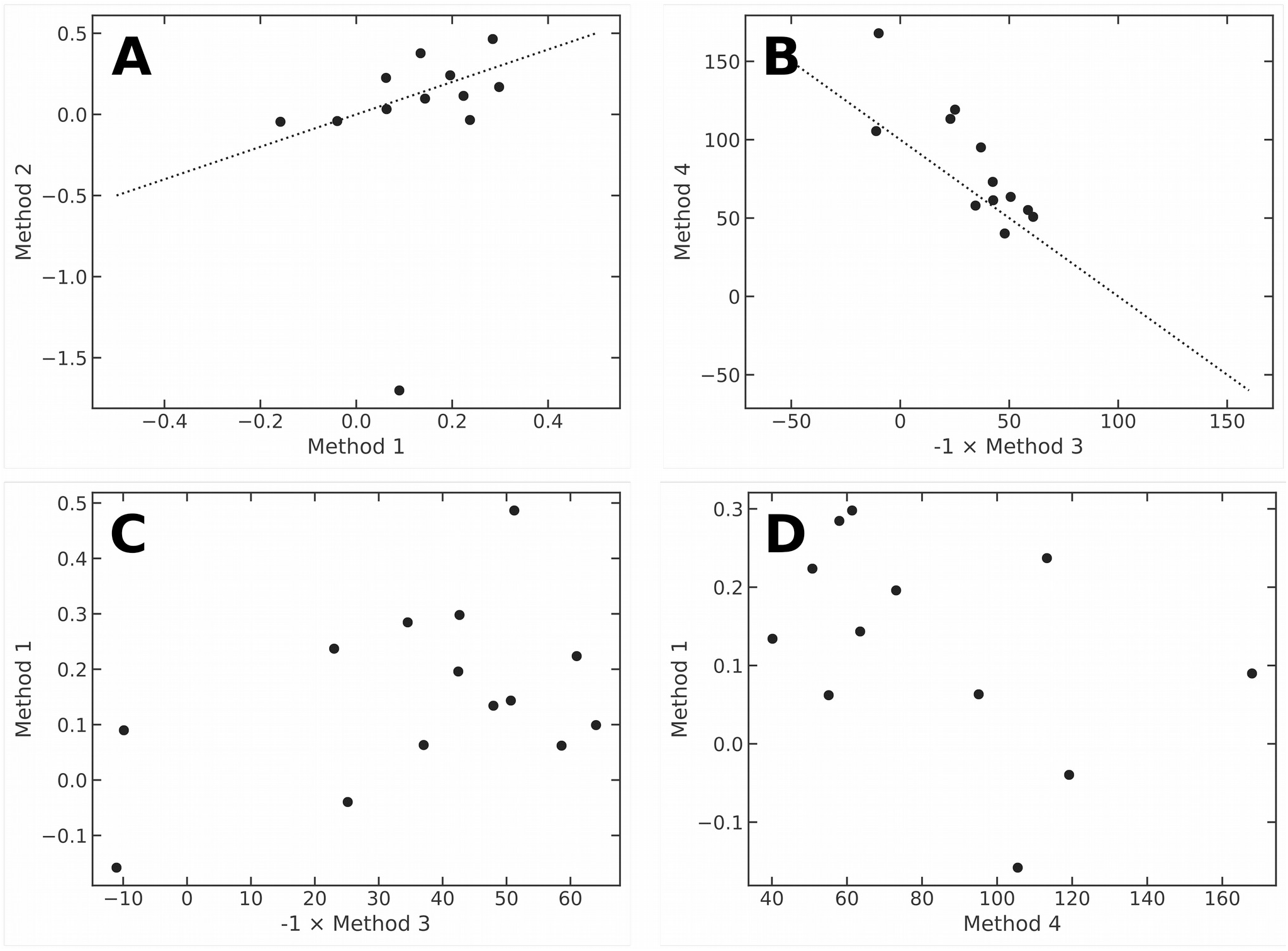
Comparison of Methods 1, 2, 3, and 4 for estimating ecosystem resistance to drought for the 14 herbaceous systems (see Table 1 for definitions). Panel **A)** compares two ‘slope-based’ methods, panel **B)** compares to ‘percent-based’ methods, while panels **C)** and **D)** compare a ‘slope-based’ method (Method 1) to both ‘percent-based’ methods. The dashed line in panels **A)** and **B)** are the 1:1 line of perfect correspondance.

**Figure 4.**
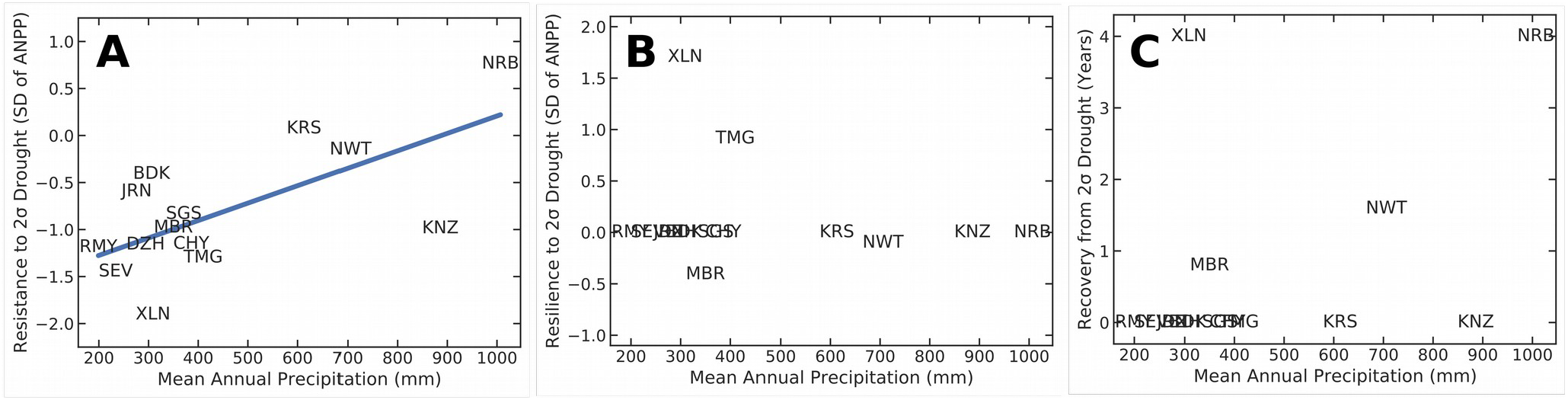
Relationship between precipitation and A) resistance, B) resilience, and C) recovery following a 2σ drought for 14 herbaceous systems.

Using the IRF method revealed that primary production at the majority of sites (71%) was best described by an ARX(0) model, indicative of no legacy effects (Table 4). Of the four sites exhibiting significant lag effects, three were best fit by an ARX(1) model while only one site was best described by an ARX(2) model (Table 4). Resistance to a 2σ decline in precipitation varied among sites from a minimum of −2.0 SD decrease in ANPP at XLN to a maximum of 0.5 SD increase in ANPP at NRB (Figure 3A). Indeed, a significant positive relationship between drought resistance and mean annual precipitation (*p* = 0.008) indicated that drier herbaceous sites were generally less resistant than mesic systems. Yet the relationship was not strong (*R*^2^ = 0.41); even dry sites varied significantly in drought resistance. For example, JRN possesses roughly the same mean annual precipitation as XLN, yet JRN was 68% more resistant to a 2σ reduction in rainfall than XLN (Fig. 3A). Such variability in drought resistance among sites of similar precipitation has been previously reported (Huxman et al. 2004) and could derive from differences in species composition, rainfall patterns (*e.g.* monsoonal, Mediterranean, etc.), or management history among sites. Relatively few sites demonstrated lag effects, such that most sites did not exhibit either resilience or recovery (Fig. 3B,C). There was no relationship between mean annual precipitation and either the strength of resilience/recovery or the probability of a legacy effect (Fig. 3B,C).

**Table 4.** ΔBIC values of ARX(p) models for 14 grassland sites used in Sala et al. (2012). Bold denotes the best model, chosen by ΔBIC < 2. In the case of multiple competing models (ΔBIC < 1), I chose the simplest model following the principle of parsimony.

| Site | ARX(<br>0) | ARX(<br>1) | ARX(<br>2) |
| --- | --- | --- | --- |
| Badkhyz, Turkmenistan<br>(BDK) | **0.0** | 2.8 | 6.3 |
| Cheyenne, Wyoming<br>(CHY) | **0.0** | 2.4 | 4.3 |
| Dzhanybek, Kazakhstan<br>(DZH) | **0.0** | 3.0 | 6.5 |
| Jornada, New Mexico<br>(JRN) | **0.0** | 3.2 | 4.8 |
| Konza Prairie, Kansas<br>(KNZ) | **0.0** | 3.3 | 6.7 |
| Kursk, Russia<br>(KRS) | **0.0** | 1.3 | 3.5 |
| Manyberries, Alberta<br>(MBR) | 4.1 | **0.0** | 2.1 |
| Nairobi, Kenya<br>(NRB) | **0.0** | 0.8 | 2.6 |
| Niwot Ridge, Colorado<br>(NWT) | 14.0 | **0.0** | 0.4 |
| Rio Mayo, Argentina<br>(RMY) | **0.1** | 2.1 | 0.0 |
| Sevilleta, New Mexico<br>(SEV) | **0.0** | 0.9 | 1.3 |
| Fort Collins, Colorado<br>(SGS) | **0.0** | 0.0 | 2.7 |
| Tumugi, China<br>(TMG) | 1.6 | **0.0** | 2.0 |
| Xilingol, China<br>(XLN) | 1.90 | 1.80 | **0.0** |

## Discussion

Given the expected increase in both the severity and intensity of extreme stress events as climate change progresses, it is imperative that we accurately quantify how ecosystems respond to stress. Estimating ecosystem vulnerability to stress using long-term time series data is a promising approach, but ecologists have not yet coupled time series data with the appropriate statistical tools. Most current methods possess flaws that potentially bias estimates of ecosystem susceptibility to stress and potentially misidentify legacy effects. To resolve these issues, I advocate for using IRFs derived from autoregressive time series models as a single quantitative framework that can accurately estimate ecosystem resistance, resilience, and recovery from severe stress events. Impulse response functions have numerous advantages over prior techniques, including the separation of observation and process errors, standardizing drought intensity among different locations, and rigorously testing for legacy effects.

Legacy effects have previously been suggested to be widespread among terrestrial ecosystems (Sala et al. 2012). However, evidence for legacy effects is mixed. Previous studies using the same datasets here reported significant autocorrelation of primary production in only six grasslands (at a significance threshold of *p* = 0.1, Sala et al. 2012), while the information theoretic approach advocated here identified significant autocorrelation in only four grasslands, albeit at a stronger threshold for significance. These qualitatively similar patterns suggest that legacy effects may be rare. This is not to say that legacy effects do not occur; controlled experiments identified strong drought legacies in the Patagonian steppe of southern Argentina (Yahdjian and Sala 2006). Experiments have even demonstrated positive legacy effects of drought in herbaceous communities of Europe (Hofer et al. 2017). Instead, it appears that grassland systems dominated by fast-growing, perennial, herbaceous species tend to be highly resilient and recover quickly from stress (Stuart-Haëntjens et al. 2018), such that drought legacies in grasslands might not be widespread. In contrast, forests are dominated by slow-growing, woody trees and shrubs and are often much less resilient to drought than grasslands (Stuart-Haëntjens et al. 2018). It is therefore plausible that legacy effects are much more common in forests than in grasslands (Anderegg et al. 2015). Testing this hypothesis using the statistical framework proposed here represents an interesting direction of future research.

The method proposed here also yields results consistent with previous studies regarding regional and global patterns in grassland resistance to drought. Multiple experimental and observational studies reported that arid and semi-arid grasslands are most sensitive to drought stress (Huxman et al. 2004, Knapp et al. 2015a). Likewise, the 2*σ* reduction in rainfall tested here provided the same result; semi-arid grasslands experience stronger reductions in primary production relative to normal inter-annual variation than did mesic grasslands. However, patterns in resilience across climates reported here did differ from previous work. Stuart-Haëntjens et al. (2018) reported strong regional variation in grassland resilience to drought; semi-arid grasslands were significantly less resilient to drought than were mesic grasslands. In contrast, the IRF method identified relatively few instances of legacy effects, such that most grasslands were recovered immediately following stress and there was no clear trend with climate. Inconsistencies between the results of this study and Stuart-Haenjens et al. (2018) likely arise from key differences in the underlying data; this study used long-term, observational time series to calculate resilience, whereas Stuart-Haenjens et al. (2018) compiled data from experimental droughts. Applying the IRF method to more grassland sites, consisting of longer time series, could prove fruitful in assessing abiotic constraints on ecosystem resilience to drought.

One advantage of the method proposed here is the relative ease with which ARX models can be fit and IRFs calculated in common statistical programming languages. The following recommendations would prove beneficial for ecologists wishing to implement the method outlined here:

1. *Properly pre-treat data* – Data must be processed properly prior to analysis with autoregressive models. First, data must be examined for gaps, as simple ARX models proposed here do not function with non-contiguous data. Small gaps can be filled with a data imputation function (*e.g.* radial basis functions, used here). Second, data must be detrended. ARX models assume stationarity, wherein the mean and variance do not change through time. Detrending data can stabilize the mean through time, but variances must still be checked visually. Third, data should be standardized in order to facilitate comparison among sites. For example, if precipitation is not standardized, then *α* = −2 for the IRF is only a 2mm decline in rainfall, rather than an extreme 2*σ* event.
2. *Use a 2σ increase or decrease in the exogenous stressor* – If all ecologists use a 2*σ* change in the exogenous stressor, then results are perfectly comparable among studies. I chose 2*σ* because it represents an extreme event. For example, a 1*σ* decline in rainfall is the 16% quantile, whereas a 2*σ* decline in rainfall represents a drought falling in the 2% quantile (assuming a normal distribution), thereby representing an extreme stress event.
3. *Report the autoregressive order and parameter values* – Reporting the parameters enables future researchers to easily extract the IRF and calculate ecosystem stress responses under different *x*^*^. For example, ecologists could standardize all IRFs to a 2*σ* stress if variation exists in the literature, or could assess ecosystem recovery using values different from a 50% return in ecosystem function. Alternatively, future researchers could use IRFs to assess how systems respond to multiple stress events of either identical or varying magnitude.
4. *Use designated AR model fitting functions* – The ARX functions specified here could all be fit using least squares. Doing so, however, requires trimming the first two data points from all model fits because we cannot use information theory or likelihood ratio tests to compare models fit to different datasets (*e.g. n* points for ARX(0), *n*−1 points for ARX(1), *n*−2 points for ARX(2), etc.). For small datasets, the loss of two data points can substantially alter the results. For example, using OLS to fit an ARX(0) model to the RMY data without the first two data points (n = 8) results in no relationship between primary production and precipitation (*p* = 0.65) because the first two data points are the driest and wettest years. Using the full dataset (*n* = 10) yields a stronger primary production – precipitation relationship (*p* = 0.12). Common statistical languages have ARIMAX functions (R: TSA library, Python: statsmodels module) wherein the user can specify the AR order, incorporate an exogenous predictor, and utilize the full dataset.

In conclusion, IRFs provide ecologists with a quick and simple means for quantifying ecosystem responses to extreme stress, while enabling ecologists to capitalize on the increased availability of long-term, observational time series data. Ecologists can use this method to quantify the components of ecosystem stress response in a standardized way across many sites. Site-specific information on species composition, long-term climate, rainfall patterns, or any other important variable can then be used to identify the abiotic and biotic factors that dictate ecosystem stress response. For example, the brief analyses presented here suggest that dry grasslands are often more sensitive to drought than wet grasslands, but also that our understanding of differential ecosystem sensitivity to drought remains incomplete. As a result, IRFs should greatly improve our ability to predict how ecosystems will respond to the increased severity and frequency of extreme events in the future.

## Acknowledgments

I would like to thank newly minted Drs. R. Griffin-Nolan and A. Hoffman for their helpful comments on drafts of this manuscript. This work was funded by an NSF DEB award (1941390) to NPL.

## Footnotes

1 Care must be taken for matrix powers in many programming languages. In R, typing psi^2 simply squares every element of psi and is not the same as psi %*% psi, which is the matrix square of psi. In Python, the square of a matrix is calculated using the matrix_power function in the numpy.linalg module.

